# Characterization of chitin deacetylase genes in the *Diaphorina citri* genome

**DOI:** 10.1101/2020.12.22.424074

**Authors:** Sherry Miller, Teresa D. Shippy, Blessy Tamayo, Prashant S Hosmani, Mirella Flores-Gonzalez, Lukas A Mueller, Wayne B Hunter, Susan J Brown, Tom D’elia, Surya Saha

## Abstract

Chitin deacetylases (CDAs) are one of the least understood components of insect chitin metabolism. The partial deacetylation of chitin polymers appears to be important for the proper formation of higher order chitin structures, such as long fibers and bundles, that contribute to the integrity of the insect exoskeleton and other structures. Some CDAs may also play a role in bacterial defense. Here we report the characterization of four CDA genes from the Asian citrus psyllid, *Diaphorina citri,* laying the groundwork for future study of these genes. *D. citri* is the vector for *Candidatus* Liberibacter asiaticus (CLas), which is responsible for the global outbreak of Huanglongbing (citrus greening) disease. The manual annotation was done as part of a collaborative community annotation project (https://citrusgreening.org/annotation/index).

## Introduction

Chitin deacetylases (CDAs) are metalloenzymes that partially deacetylate chitin polymers [1]. CDA activity in insects was first reported in the cabbage looper *Trichoplusia ni* [2]. In *Drosophila melanogaster,* several CDAs were found to play a role in tracheal development [3,4]. More recently, Genomic and phylogenetic studies have shown that CDAs are present widely in insects and can be classified into five different groups [5,6]. Most holometabolous insects have at least one representative of each of the five CDA groups, while the hemimetabolous insects that have been examined lack group II and group V genes [7]. The exact role of insect CDAs is not well understood, but they may play a role in organization of chitin molecules into higher order structures [8]. Loss of function experiments indicate that some CDAs play an essential role in growth and development, making them a potential target for insect pest control [6,9–12]. Here we describe the chitin deacetylase gene family in the Asian citrus psyllid, *Diaphorina citri. D. citri* is the vector for *Candidatus* Liberibacter asiaticus (CLas), which is responsible for the global outbreak of Huanglongbing (citrus greening) disease. We identified four chitin deacetylase genes in the *D. citri* v3 genome, three of which have multiple isoforms. As in other hemipterans, only groups I, III and IV are represented [6].

## Results

Chitin deacetylase genes in the *D. citri* v3 genome [13] were identified and manually annotated. These genes were classified into groups following the precedents established in other insects [5,6].

### Group I chitin deacetylases

Most insects have two group I genes named *CDA1* and *CDA2* (Table 1). The proteins encoded by these genes have an N-terminal chitin-binding domain (ChBD), a low-density lipoprotein receptor class A domain (LDLa), and a deacetylase catalytic domain [5]. RNAi of group I CDAs in a variety of insects suggests that loss of function of CDA1 or CDA2 can result in lethality and therefore these genes could be potential targets for pest control methods [6,8–12,14]. Recent experiments in *Tribolium* suggest that TcCDA1 and TcCDA2 are required for organization of chitin into longer fibers that are important for cuticular strength [8].

**Table 1.** *D. citri* gene numbers were determined based on annotation of the *D. citri* genome v3. All other ortholog numbers were obtained from published sources [5,6,29–37]. An asterisk (*) indicates that isoforms have been found for at least one member of the group in that organism.

|  | Group I | Group II | Group III | Group IV | Group V | Total |
| --- | --- | --- | --- | --- | --- | --- |
| *D. melanogaster* | 2* | 1 | 1 | 1* | 1 | 6 |
| *A. gambiae* | 2* | 1 | 1 | 1 | 0 | 5 |
| *T. castaneum* | 2* | 1 | 1 | 1* | 4 | 9 |
| *B. mori* | 2* | 1 | 1 | 1 | 3 | 8 |
| *A. mellifera* | 2* | 1 | 1 | 1* | 0 | 5 |
| *N. vitripennis* | 2 | 1 | 1 | 1 | 0 | 5 |
| *R. prolixus* | 2 | 0 | 1 | 1 | 0 | 4 |
| *A. pisum* | 2 | 0 | 1 | 1 | 0 | 4 |
| *N. lugens* | 2 | 0 | 1 | 1 | 0 | 4 |
| *D. citri* | 2* | 0 | 1 | 1* | 0 | 4 |

As expected, we identified two group I genes in *D. citri,* which we named *CDA1* and *CDA2.* Both genes encode proteins with the typical group I domain structure (Figure 1). We identified two isoforms each for *D. citri CDA1* and *CDA2* (Table 2). *CDA2* has previously been shown to have multiple isoforms in several holometabolous insect species, with the transcripts differing only in the use of one alternative exon [5,11,14]. This gene structure is conserved in *D. citri CDA2* with alternate exons 3a and 3b. The two *D. citri* CDA1 isoforms differ in the presence or absence of a 24 bp exon upstream of the last exon. Expression data from RNA-Seq datasets available through the Citrusgreening Expression Network (CGEN) [15] suggest that, in general, expression of CDA1 and CDA2 is higher in nymphs and eggs than in adults (Fig. 2A).

**Figure 1.**
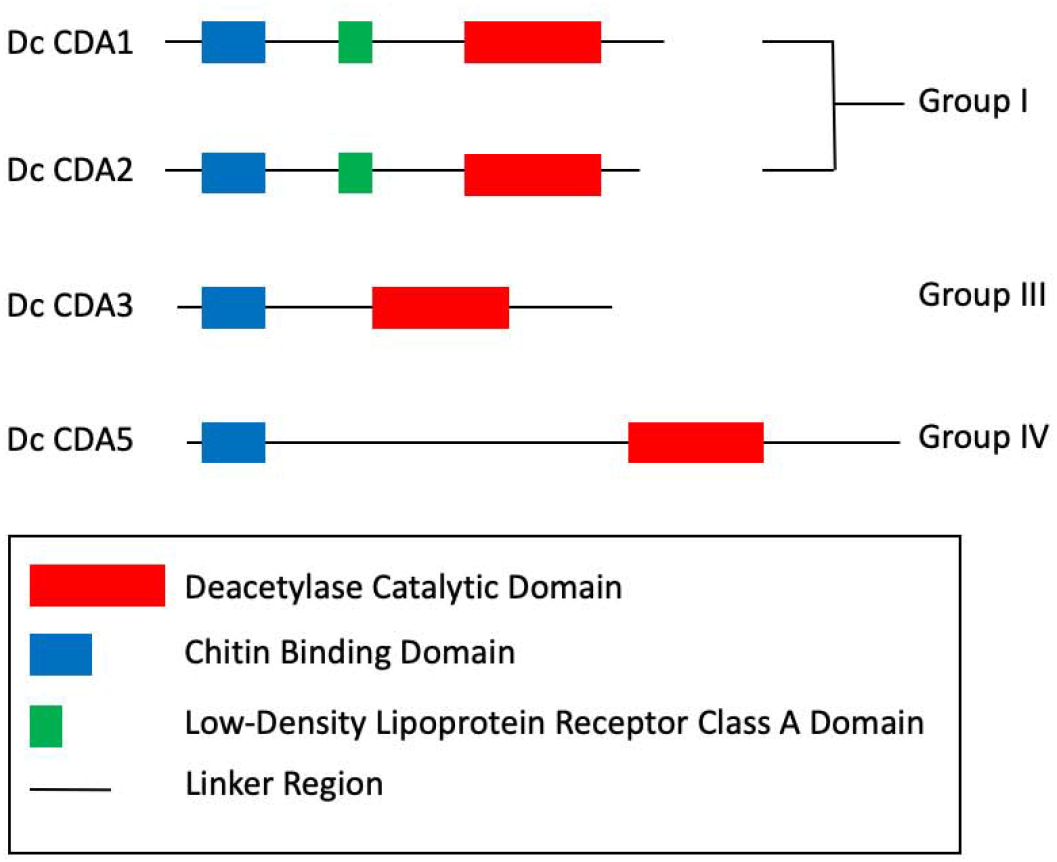
Chitin Deacetylase Domain Organization in *D. citri.* Chitin deacetylases are categorized by group based on phylogenetic analysis, sequence similarity, and domain organization. *D. citri* domain analysis was performed using InterPro [28]. CDA5 is represented by the protein encoded by the *de* novo-assembled transcript MCOT06229.1.CO [17] because a small portion of the *CDA5* gene is missing from the genome.

**Figure 2.**
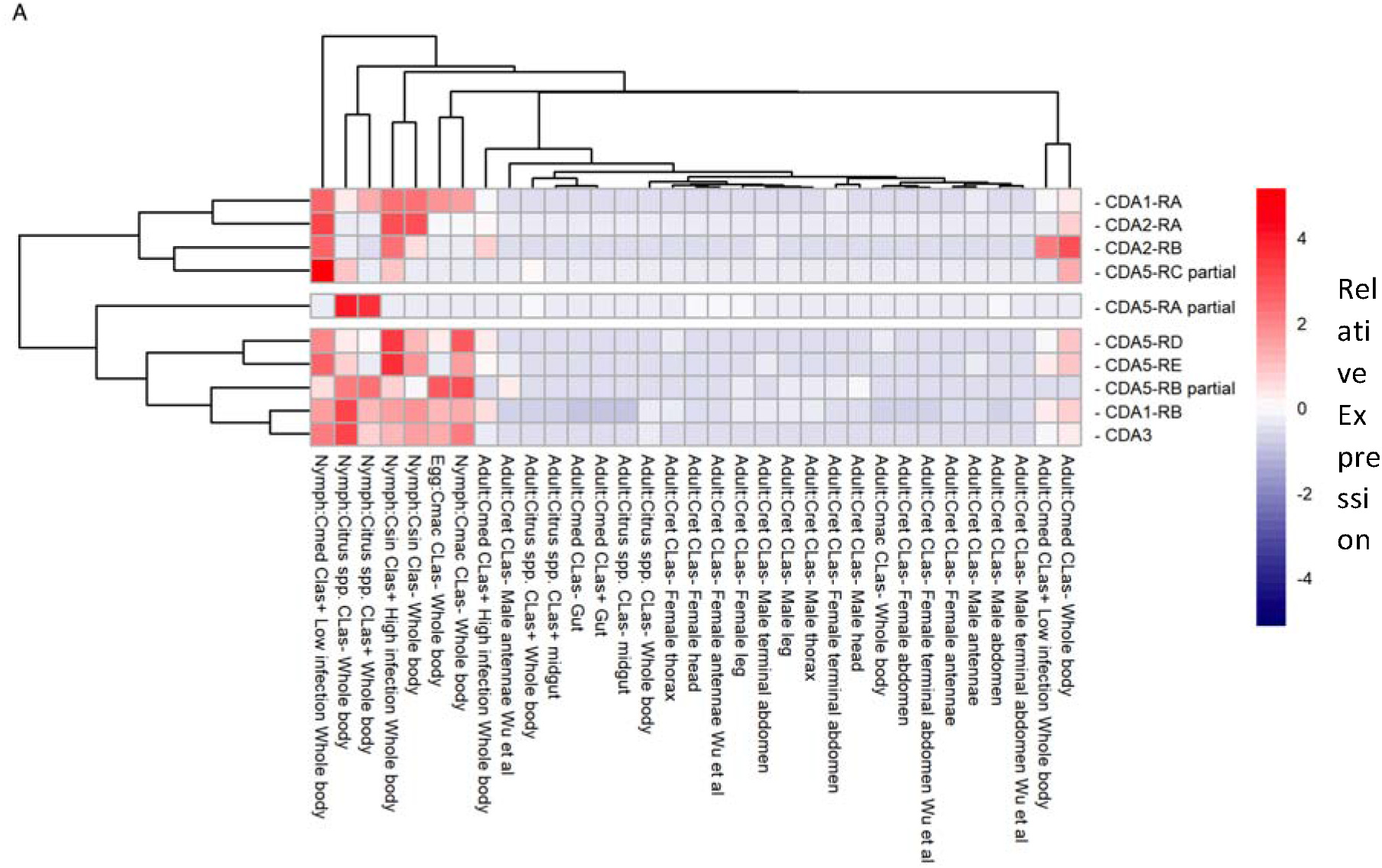
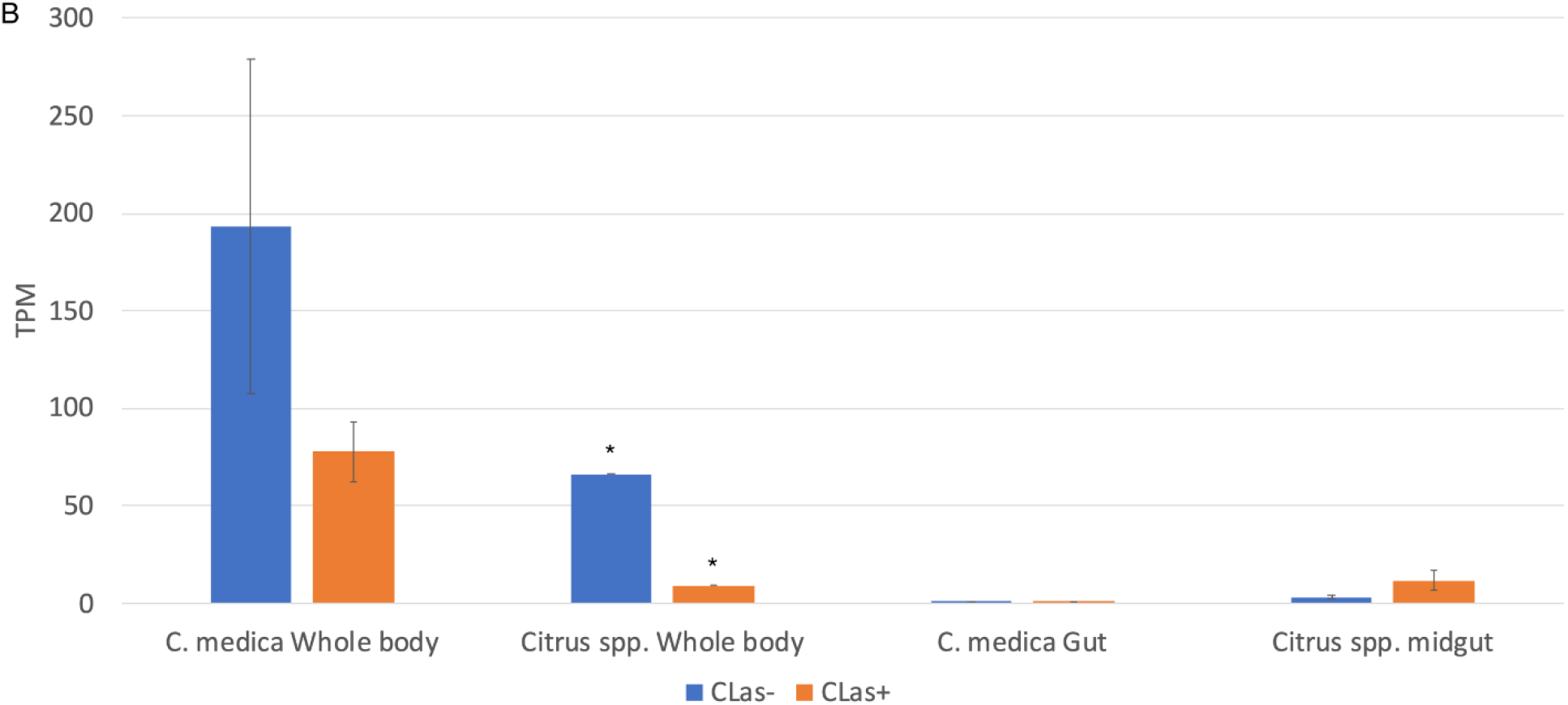
(A) Heatmap displaying relative expression levels of all annotated chitin deacetylase genes in RNA-Seq datasets from various life stages, tissues and CLas infection states. (B) Expression levels (TPM) of CDA3 (Dcitr02g03950.1.1) in tissues from CLas÷ and CLas-psyllids fed on two different types of citrus plants. Standard error bars are shown for all expression values except those marked with an asterisk (*), which had only one replicate. Expression levels were obtained from the Citrusgreening Expression Network [15].

**Table 2.**
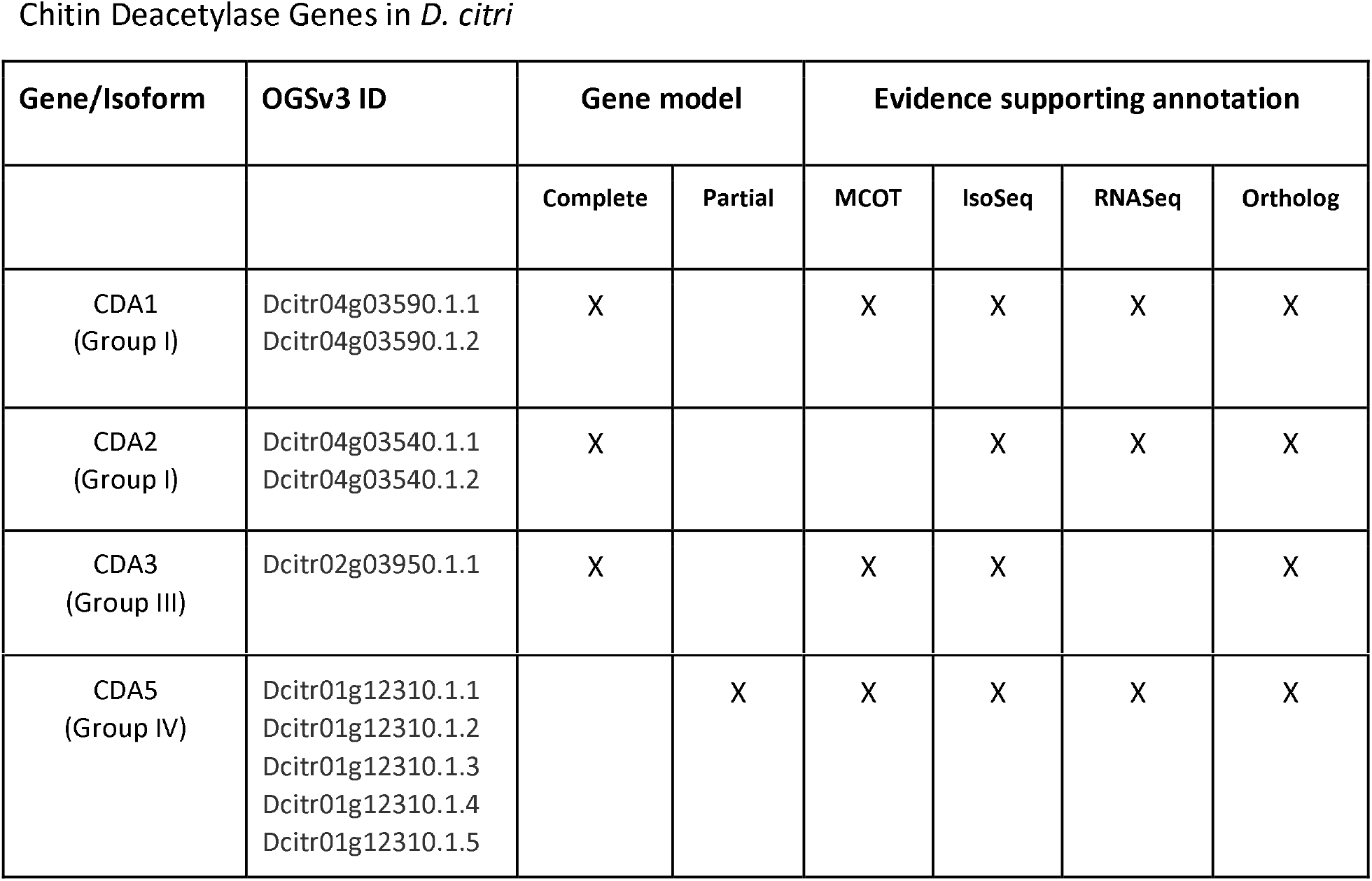
Each manually annotated gene has been assigned an OGSv3.0 gene identifier. Each gene has been termed partial or complete based on available evidence. Evidence types used for manual annotation of each gene are also indicated. More information on these evidence sources is available in [22].

In *Drosophila* and *Tribolium,* the *CDA1* and *CDA2* orthologs are adjacent to one another in the genome [3,5] on chromosome 3 and 5, respectively. The conserved clustering of these genes suggests there may be evolutionary constraint on their physical location. We found that the *D. citri CDA1* and *CDA2* orthologs are also adjacent to one another on chromosome 4. In the *D. citri* v3 genome these genes are separated by approximately 50 kb, although this distance appears to be inflated by falsely duplicated fragments of both genes in this assembly.

### Group III chitin deacetylases

We identified one group III CDA in the *D. citri* v3 genome (Table 1). This gene has been previously described and was named *CDA3* because of its orthology to *Nilaparvata lugens CDA3*[16]. Like group III CDAs in other insects, *D. citri* CDA3 contains a ChBD and catalytic domain but lacks the LDLa domain found in group I CDAs (Figure 1). Due to improvements in the genome assembly, our curated CDA3 model from genome v3 has additional 3’ sequence compared to the previously reported model, which was based on genome v1.1 [16,17]. The resulting predicted protein is almost 50 amino acids longer, with additional conserved sequence at the C-terminus.

Yu et al. [16] reported that RNAi knockdown of *CDA3* had no effect on molting or wing development. Instead, their results implicated *CDA3* in the *D. citri* bacterial immune response. Recombinant CDA3 showed antibacterial activity against gram-positive bacteria, but had no effect on gram-negative bacteria. Moreover, injection of either *Escherichia coli* (gram-negative) or *Staphylococcus aureus* (gram-positive) bacteria into *D. citri* increased *CDA3* expression in the midgut and decreased its expression in the fat body, although the timeline of these effects is not certain. To determine whether infection by CLas, a gram-negative bacterium, might also affect *CDA3* expression, we used CGEN [15] to compare expression of *D. citri CDA3* in RNA-Seq datasets from CLas+ and CLas-guts [18], midguts [19] and whole bodies ([20] and NCBI BioProject PRJNA609978).

*CDA3* expression in was lower in CLas+ versus CLas-whole body tissue in data from two different RNA-Seq experiments (Figure 2B). Expression of *CDA3* in midgut and gut tissues was very low in all samples but there was a slight increase in CLas+ versus CLas-midgut expression (Figure 2B). While the significance of these expression differences is not clear, they may warrant further investigation.

### Group IV chitin deacetylases

Most insects examined to date have one group IV CDA, typically called *CDA5 (CDA4* in *N. lugens)* (Table 1). CDA5 has been shown to have multiple isoforms in *Tribolium* and *Drosophila* [5]. Consistent with these observations, we identified and annotated five different isoforms of CDA5 in *D. citri* (Table 2). Unfortunately, the annotated models are missing a small amount of 3’ sequence due to genome assembly issues. However, we identified a *de-novo* assembled transcript (MCOT06229.1.CO) that appears to encode the full-length protein (Figure 1). The missing genome sequence does not affect the conserved function domains of CDA5. Four of the five transcript isoforms encode proteins containing both an N-terminal ChBD and a C-terminal catalytic domain, as seen in other insect CDA5 orthologs. The remaining isoform (CDA5-RB) differs at the 5’ end and apparently lacks a ChBD-encoding region.

### Other chitin deacetylase groups

We did not find any group II or group V CDAs in the *D. citri* v3 genome (Figure 3). To our knowledge CDAs from these groups have not been found in any hemipteran insects examined to date [6], so their absence in *D. citri* was expected.

**Figure 3.**
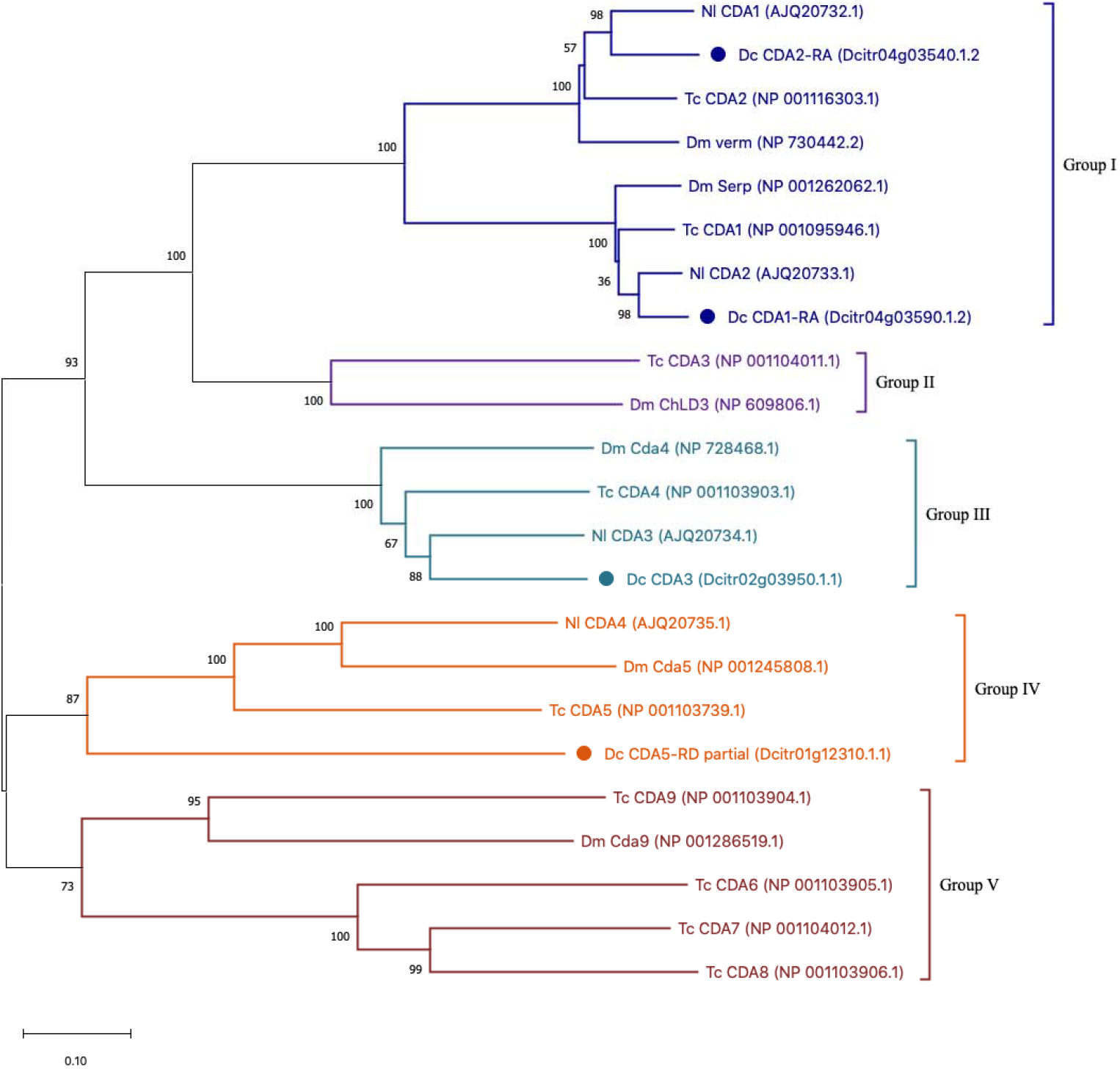
Phylogenetic tree of chitin deacetylase family members. ClustalW was used to perform the multiple sequence alignment. The tree was constructed with MEGA X software using neighbor-joining analysis with 100 bootstrap replications. *Drosophila melanogaster* (Dm), *Tribolium castnaeum* (Tc), *Nilaparvata lugens* (Nl), and *Diaphorina citri* (Dc). *D. citri* protein branches are marked with a circle. Colors delineate established chitin deacetylase groups.

## Conclusion

Chitin is a polysaccharide found primarily in invertebrates, where it is required for the exoskeleton and other structures. Chitin deacetylases modify these molecules and are believed to play a role in the assembly of higher order chitin structures, although their precise role in insects is not well characterized. At least some of the members of this gene family seem to be required for normal development in many insects. The essentiality and limited phylogenetic distribution of these genes makes them attractive targets for pest control via gene knockdown methods such as RNAi. As part of a community project to annotate the genome of *D. citri,* the vector of Huanglongbing (citrus greening disease), we identified and annotated four chitin deacetylase genes in the *D. citri* v3 genome. Three of these genes have multiple transcriptional isoforms, so having correctly annotated transcripts will be particularly valuable for any future experiments targeting these genes.

## Materials and Methods

Chitin deacetylase genes in *D. citri* genome v3 [13] were identified by BLAST search of *D. citri* sequences with chitin deacetylase orthologs from other insects. Orthology was confirmed by reciprocal BLAST of the NCBI non-redundant protein database. Genes were manually annotated in Apollo 2.1.0 [21] using available evidence, including RNA-Seq reads, IsoSeq transcripts and *de*novo-assembled transcripts. A more detailed annotation protocol is available at protocols.io [22]. Multiple alignments were performed using MUSCLE [23] and phylogenetic trees were constructed in MEGA7 [24] or MEGA X [25]. Table 3 contains a list of orthologs used in the phylogenetic analysis. Expression data from CGEN [18] was visualized using the pheatmap package of R [26,27] or Microscoft Excel.

**Table 3.**
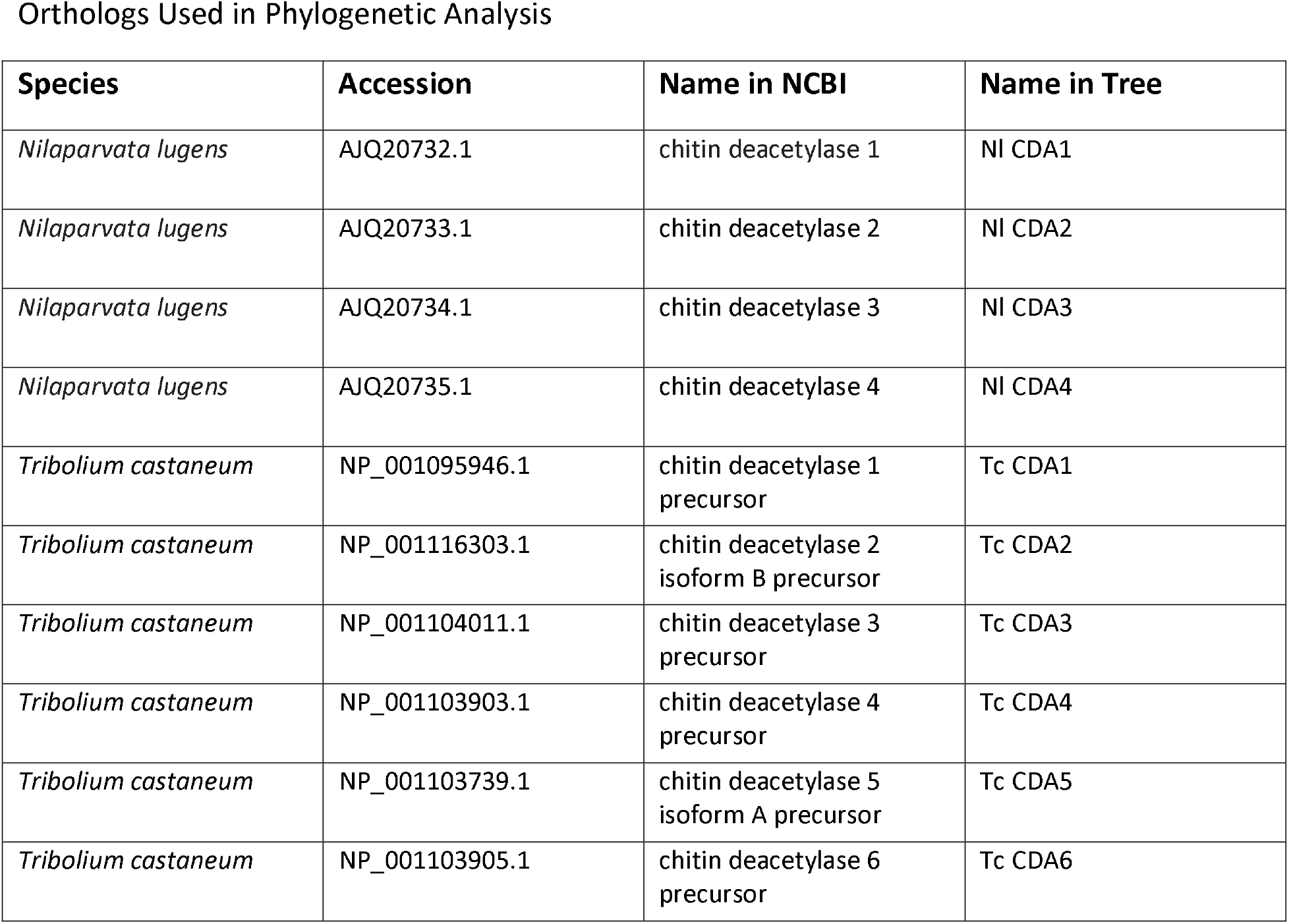
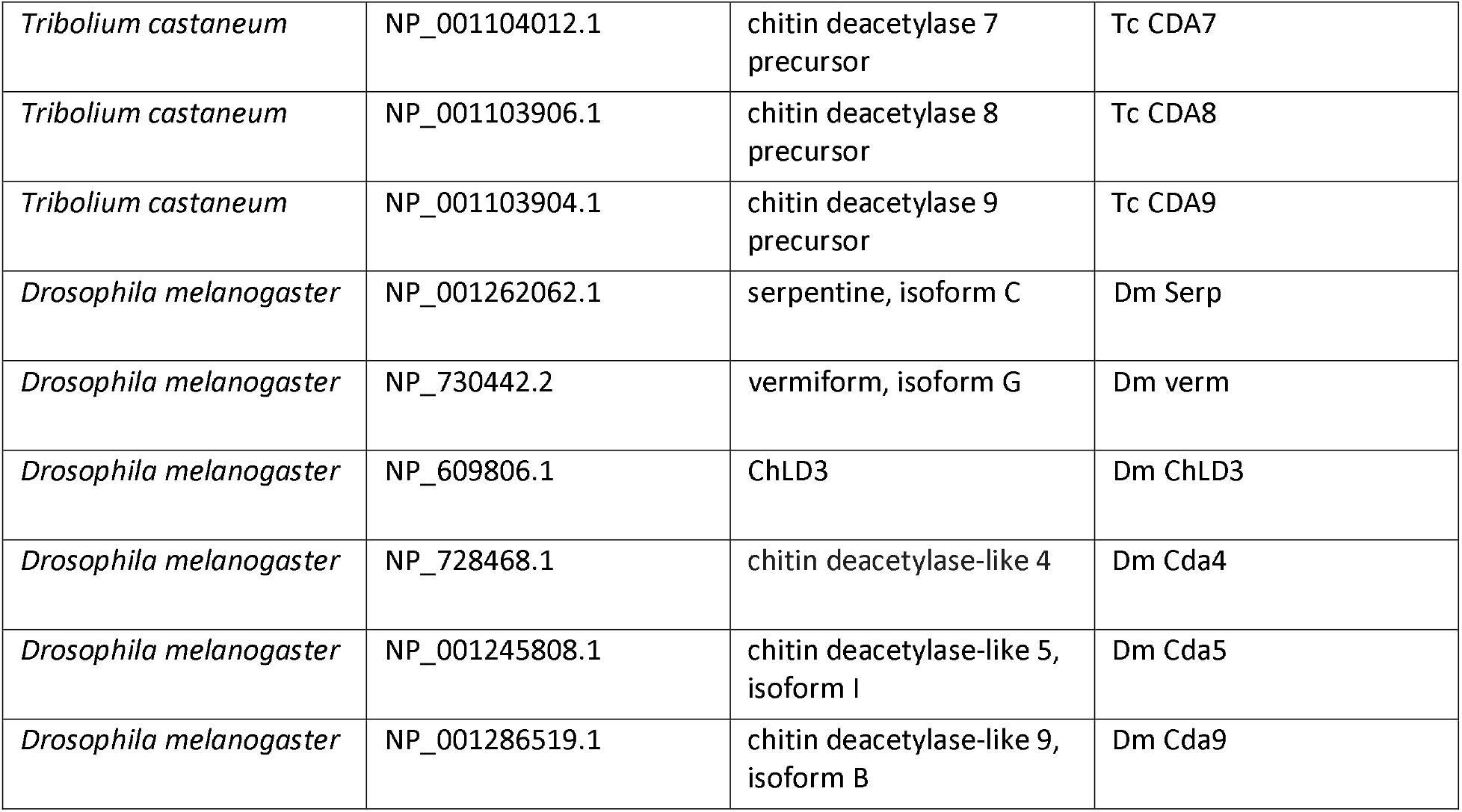
Species, accession number, full name and abbreviated name are provided for all orthologs used in phylogenetic analysis.

## Acknowledgements

We thank Dr. Josh Benoit for assistance with data visualization. This research was funded by USDA-NIFA grant 2015-70016-23028 and an Institutional Development Award (IDeA) from the National Institute of General Medical Sciences of the National Institutes of Health under grant number P20GM103418.

## Author Contributions

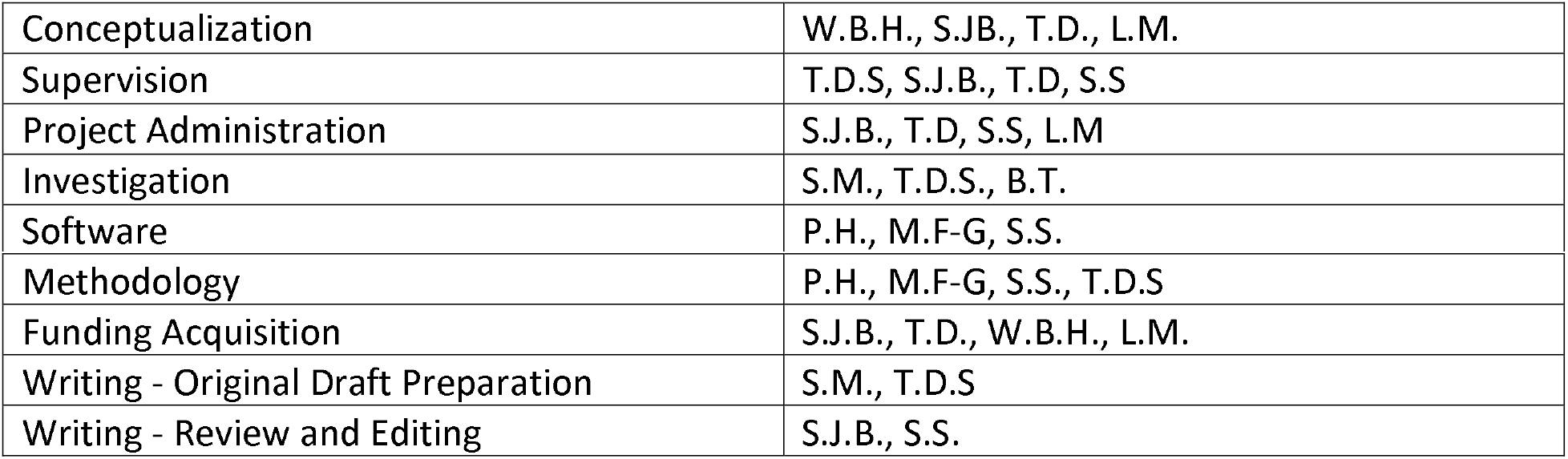

## Notes

https://citrusgreening.org/organism/Diaphorina_citri/genome

https://citrusgreening.org/annotation/index

### Competing Interest Statement

The authors have declared no competing interest.

